# Qualitative analyses of polishing and pre-coating FIB milled crystals for MicroED

**DOI:** 10.1101/613042

**Authors:** Michael W. Martynowycz, Wei Zhao, Johan Hattne, Grant J. Jensen, Tamir Gonen

## Abstract

Microcrystal electron diffraction (MicroED) leverages the strong interaction between matter and electrons to determine protein structures from vanishingly small crystals. This strong interaction limits the thickness of crystals that can be investigated by MicroED, mainly due to absorption. Recent studies have demonstrated that focused ion beam (FIB) can thin even very large crystals into ideal sized lamellae however it is not clear how to best apply FIB-milling for MicroED. Here, The effects of polishing the lamellae, whereby the last few nanometers are milled away using a low-current gallium beam, are explored in both platinum pre-coated and uncoated samples. Our results suggest that pre-coating samples with a thin layer of platinum followed by polishing the crystal surfaces prior to data collection consistently led to superior results as indicated by higher signal/noise ratio, higher resolution and better refinement statistics. This study lays the foundation for routine and reproducible methodology for sample preparation in MicroED.

## Introduction

Focused ion beam (FIB) milling is a mature technique for shaping samples in materials science, and has recently gained popularity for preparing biological samples for investigation in a transmission electron microscope (TEM) by cryoEM (Marko et al., 2007; Rigort et al., 2012; Wirth, 2009). Many biological samples such as cells and tissues are too thick, and would be inaccessible to TEM investigation without sectioning or milling them down to sub-micron thicknesses (Marko et al., 2007). When milling, FIB rasters back and forth across the sample to remove unwanted material. In this way, the sample and surrounding material is exposed. High-speed ions deposit some fraction of their energy into the sample due to inelastic scattering. This exposure builds up while the crystal is milled, likely damaging the specimen either directly from ion bombardment or through secondary effects. Furthermore, the milling process is typically conducted at grazing incidence angles ∼18°, where large areas of the grid around the sample are also exposed to high-energy gallium ions.

Metal coating prior to FIB milling has become standard practice in FIB/SEM applications to protect the sample from the ion beam. Platinum is one of the typical materials used as a coating material because of its high conductivity and contrast in electron microscopy. This additional layer protects the sample from high-energy ions during imaging with either the electron or ion beam (Rigort et al., 2012), improves contrast for biological materials, and reduces charging effects caused by the ion beams during imaging or milling (Munroe, 2009). Platinum coating may also increase the long term stability of the grid and help preserve the specimen for future investigations.

FIB Milling crystals has recently gained attention as a method for preparing samples for microcrystal electron diffraction (MicroED) experiments. MicroED is an electron cryo-microscopy (cryo-EM) technique that uses a TEM to collect electron diffraction data from nanocrystals at cryogenic temperatures (Shi et al., 2013) under continuous rotation (Nannenga et al., 2014). Electrons at commonly used accelerating voltages (80-300kV) can only penetrate samples less than ∼1μm thickness due to the strong interaction between matter and the electron beam. Thin, uniform crystals are ideal for MicroED experiments, as this both reduces noise and eliminates the uncertainty about sample thickness (Grimm et al., 1996; Hattne et al., 2015; Martynowycz et al., 2018). Machined crystals have been shown to retain their high-resolution information, allowing structure determination at near-atomic resolution (Duyvesteyn et al., 2018; Li et al., 2018; Martynowycz et al., 2018; Zhou et al., 2019).

The first studies on FIB milled crystals were all conducted using different experimental approaches, different samples, various detectors, and refinement software. It is thus unclear which FIB milling approach results in the highest quality data. For example, Duyvesteyn *et al.* investigated milled lysozyme crystals that had been platinum-coated (Duyvesteyn et al., 2018). Their data were collected as stills from different orientations of a single crystal, and diffraction was observed to 1.9Å. They were able to achieve an overall completeness of 39% using a total exposure of ∼25e^−^Å^−2^, and used rigid body refinement to achieve reasonable structural statistics with an R_work_ and R_free_ of 29.1 and 28.3%, respectively. Li *et al.* also pre-coated their grids of lysozyme crystals with a platinum layer (Li et al., 2018) and similarly collected diffraction data as a sequence of still images with a total exposure of ∼9e^−^Å^−2^, or about threefold lower exposure than that of Duyvesteyn *et al*. Data from 7 crystals were merged to solve the structure of lysozyme to 2.5Å with an overall completeness of 94.0%. However, it appears that the model was not fully refined and a rather high R_free_ value (∼40%) was reported.

In our previous MicroED FIB milling study, we collected a continuous rotation dataset from a single proteinase K crystal to a completeness of 88% at a resolution of 2.75Å, but the samples were not pre-coated with platinum (Martynowycz et al., 2018). Proteinase K was solved by molecular replacement, and refined to final R_work_ and R_free_ values of 22 and 26%, respectively, using a much lower total exposure of 4e^−^Å^−2^. Later, Zhou *et al.* investigated lysozyme and proteinase K using an oscillation method similar to continuous rotation to solve the structures to 1.73Å and 1.50Å, respectively (Zhou et al., 2019). Both of these structures were also pre-coated with a platinum layer, and some optimization of sample thickness was conducted. The two structures were ultimately solved using data merged using four crystals each with overall completeness of 89% (lysozyme) and 91% (proteinase k), with corresponding R_work_/R_free_ values of 23/25% and 19/23%. The total exposure in this study was ∼4.5e^−^Å^−2^. Importantly, these examples used different milling procedures and data collection strategies, and it is unclear whether the resolutions and structure quality of these experiments are limited by sample, sample preparation, FIB milling protocols, radiation damage, data collection or processing.

Here we performed a systematic study to evaluate the best practices for sample preparation for MicroED. Since data from each crystal was collected in the same way we were able to perform a qualitative analyses of FIB milling procedures and evaluate how pre-coating grids with a thin layer of platinum and crystal polishing affect data quality.

## Results

### Experimental setup

Proteinase K crystals, all from the same crystallization drop, were used to create two cryoEM grids under identical blotting and freezing conditions. One of the grids was coated with a thin layer of platinum while the other was not. These grids were then clipped and loaded on to a FIB-SEM instrument where the crystals were subsequently milled into thin ∼200nm lamellae for MicroED investigation.

Crystals were selected for milling using identical selection criteria. Namely, crystals were approximately similar in size, and were at least 20μm away from the nearest grid bar, not within 10μm of another crystal, and not along the milling path of another selected crystal to prevent additional exposure to the FIB. These stringent constraints reduced the number of potential crystals to ∼10 crystals per grid. These crystals were milled to ∼200nm in thickness either with or without platinum coating. We ultimately collected data from 8 crystal lamellae of proteinase K without platinum coating and from 5 lamellae with platinum coating. Grids were transferred to a TEM for MicroED data collection as previously described (Martynowycz et al., 2019). Continuous rotation data were collected from each lamella, covering a total angular range of 60° per crystal and using a total exposure of 3e^−^Å^−2^. All data were collected and processed using the same protocol, and, because the completeness from each lamella was sufficiently high, a structure could be determined from each well-diffracting lamella.

### Effects of crystal polishing

For both the pre-coated and uncoated grids, we milled each crystal into a thin lamellae of approximately 300nm thick. This initial milling followed our previously used protocol of slowly stepping down the beam current from >300pA to 30pA. For two lamellae from each grid, we continued to mill an additional ∼50nm from each side of the lamellae to reach the final thickness of 200nm. For all of the rest of the crystals, we milled away these last 50nm using an ultra-low beam current of 10pA to create 200nm thick lamellae, a process typically referred to as polishing. This step may prevent morphological pathologies in the crystals due to the damaging nature of the high-current gallium beam, but greatly increases the amount of time it takes to reach the final lamellae thickness.

Compared to polished samples, the unpolished crystals, whether coated or not, exhibit visible pathologies when imaged (**Figure 1, lower left**), most notably clear striations parallel to the direction of the ion beam. Crystals that were neither coated with a platinum layer nor polished resulted in either no diffraction at all or lamellae that had broken away from the grid entirely (**Figure 1**). For the pre-coated samples, some diffraction spots were split or appeared smeared similarly to diffraction from multiple or even mosaic crystals. However, these reflections were ultimately integrated, with a noticeable decrease in resolution and worse merging statistics (**Table 1**). Internal consistency measures such as merging R factors and CC_1/2_ were also worse, but that may again be a consequence of the decreased resolution.

**Table 1.** Statistics and analysis for MicroED data collected under different preparation conditions.

| Table1. MicroED data from single lamellae under different preparation conditions |  |  |  |  |  |  |  |  |  |  |
| --- | --- | --- | --- | --- | --- | --- | --- | --- | --- | --- |
| Parameters |  | Polished and uncoated |  |  |  | Unpolished and coated |  | Polished and coated |  |  |
| Crystal # |  | 2 | 5 | 7 | 8 | 9 | 12 | 10 | 11 | 13 |
| Data collection | Accelerating voltage (kV) | 200 |  |  |  |  |  |  |  |  |
|  | Electron source | Field emission gun |  |  |  |  |  |  |  |  |
| | Total exposure ($e^{-}\text{\AA}^{-2}$) | 3 | | | | | | | | |
| | Wavelength ($\text{\AA}$) | 0.0251 | | | | | | | | |
|  | No. of crystals (#) | 1 |  |  |  |  |  |  |  |  |
|  | Microscope | Thermo Fischer Talos Arctica |  |  |  |  |  |  |  |  |
|  | Camera | CetaD |  |  |  |  |  |  |  |  |
| | Rotation rate ($^{\circ}\text{s}^{-1}$) | 0.2 | | | | | | | | |
| Data analysis | Resolution range ($\text{\AA}$) | 35.25 - 2.17<br>(2.248 - 2.17) | 41.63 - 2.176<br>(2.254 - 2.176) | 43.47 - 2.59<br>(2.683 - 2.59) | 35.41 - 2.17<br>(2.248 - 2.17) | 30.23 - 2.08<br>(2.154 - 2.08) | 33.66 - 2.07<br>(2.144 - 2.07) | 28.39 - 1.91<br>(1.978 - 1.91) | 30.07 - 1.85<br>(1.916 - 1.85) | 33.54 - 1.79<br>(1.854 - 1.79) |
| | Unit cell<br>(a=b,c) ($\text{\AA}$)<br>($\alpha=\beta=\gamma$) ($^{\circ}$) | 67.3 104.9<br>90 | 67.6 105.7<br>90 | 67.5 105.2<br>90 | 67.5 105.6<br>90 | 67.6 104.5<br>90 | 67.3 105.1<br>90 | 67.5 105.9<br>90 | 67.2 107.0<br>90 | 67.1 107.0<br>90 |
|  | Space group | P 4<sub>3</sub> 2<sub>1</sub> 2 |  |  |  |  |  |  |  |  |
|  | Total reflections (#) | 63495 (6293) | 63830 (6496) | 34844 (3727) | 61959 (6373) | 70454 (7196) | 60628 (5059) | 88818 (9016) | 99035 (10382) | 100799 (8463) |
|  | Multiplicity | 6.3 (6.4) | 4.8 (5.0) | 4.7 (5.0) | 5.3 (5.6) | 5.1 (5.3) | 6.9 (5.9) | 5.0 (5.2) | 5.9 (6.3) | 6.8 (5.8) |
|  | Completeness (%) | 71.26 | 94.46 | 92.89 | 84.79 | 90.80 (90.40) | 57.59 (56.90) | 90.41 (90.83) | 76.93 (77.64) | 62.36 (60.91) |
| | Mean I/σI | 3.38 (0.51) | 2.68 (0.51) | 2.49 (0.89) | 3.37 (0.81) | 3.65 (0.75) | 4.04 (0.93) | 4.16 (0.60) | 4.87 (1.00) | 5.95 (0.72) |
| | Wilson B-factor ($\text{\AA}^2$) | 32.5 | 32.46 | 25.8 | 30.08 | 29.74 | 33.21 | 26.74 | 24.47 | 25.71 |
|  | R<sub>pim</sub> | 0.167 | 0.222 | 0.275 | 0.167 | 0.156 | 0.109 | 0.128 | 0.107 | 0.078 |
|  | CC<sub>1/2</sub> | 0.966 | 0.956 | 0.898 | 0.956 | 0.973 | 0.975 | 0.984 | 0.987 | 0.991 |
|  | R<sub>work</sub> | 0.220 | 0.223 | 0.256 | 0.229 | 0.223 | 0.216 | 0.197 | 0.198 | 0.198 |
|  | R<sub>free</sub> | 0.265 | 0.265 | 0.310 | 0.284 | 0.258 | 0.269 | 0.234 | 0.234 | 0.236 |

**Figure 1.**
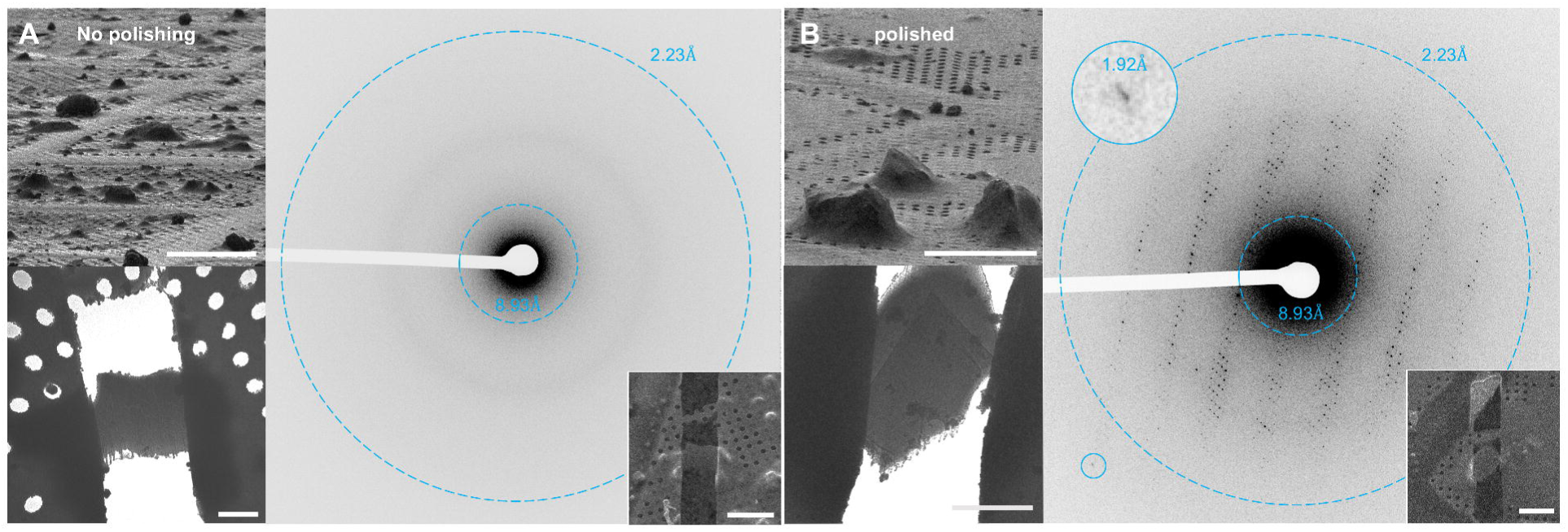
Uncoated crystals with or without polishing. The machining process of uncoated crystals without (**A**) and with (**B**) a final polishing step. Focused ion beam images of the selected crystals before milling them down to 200nm thick lamellae (Top left). Resulting diffraction patterns from the selected lamellae (Right). Images of the final, prepared lamellae (right, inset). Clear striations and outline of the crystalline portions of these lamellae are visible in the over-focused diffraction images (bottom left). Scale bars 50μm top left, 5μm bottom left, 20μm right inset.

### Proteinase K crystals milled without platinum pre-coating

We were able to identify 8 crystals that met our selection criterion from the uncoated grid. These crystals appeared to be randomly oriented upon the carbon support film. Each of these crystals was milled to a thickness of ∼200nm as detailed in **Methods**. Of these, 2 were not polished and 6 were polished. These grids were transferred to a TEM, and continuous rotation MicroED data were collected from each crystalline lamella (**Figure 1**). Of these 8 lamellae, 4 yielded clear diffraction spots, three showed no diffraction or were detached from their support, and one diffracted to ∼8Å and could not be indexed to the correct point group (**Figure 1**), even when enforcing the known unit cell. All four integrated datasets were sufficiently complete to allow a structure to be determined by molecular replacement using the search model 6CL7. Structures were refined in phenix.refine using electron scattering factors, automatic solvent picking, and without any manual intervention such as modelling ions. This approach was chosen to eliminate user bias and assure proper comparisons between solved structures. A complete list of statistics are presented in **Table 1**. The differences in completeness for these crystals are expected, as our previous works have demonstrated that proteinase K tends to orient randomly on the grid (Hattne et al., 2016, 2018a; de la Cruz et al., 2017). The average maximum resolution for these crystals was 2.28Å. The averaged, final refinements of these crystals had an R_work_ of 23.2% and an R_free_ of 28.1% and showed no clear pathologies or issues in the densities. A typical uncoated crystal, final milled lamella, diffraction pattern, and solved structure are given in **Figure 1**.

### Proteinase K crystals milled after platinum pre-coating

A second grid with crystals from the same crystallization well was prepared identically. This grid was coated in a thin layer of platinum (10-50nm) prior to being loaded into the FIB/SEM instrument. Using the same criteria as the uncoated grid, we located and milled five crystals. Three of these were polished, and two unpolished as indicated. Continuous rotation MicroED data were then collected as above. All 5 crystals yielded diffraction that indexed correctly. Each crystalline lamella was solved similarly to the uncoated lamellae above, and the individual statistics for these solutions are also presented in **Table 1**. A typical platinum-coated, polished crystal with the corresponding lamella, and diffraction pattern are presented in **Figure 2,** alongside an typical uncoated, polished crystal. On average, these crystals diffracted to a resolution of 1.94Å with an R_work_ of 20.6% and an R_free_ of 24.6%. Statistics for individual platinum coated lamellae are given in **Table 1**.

**Figure 2.**
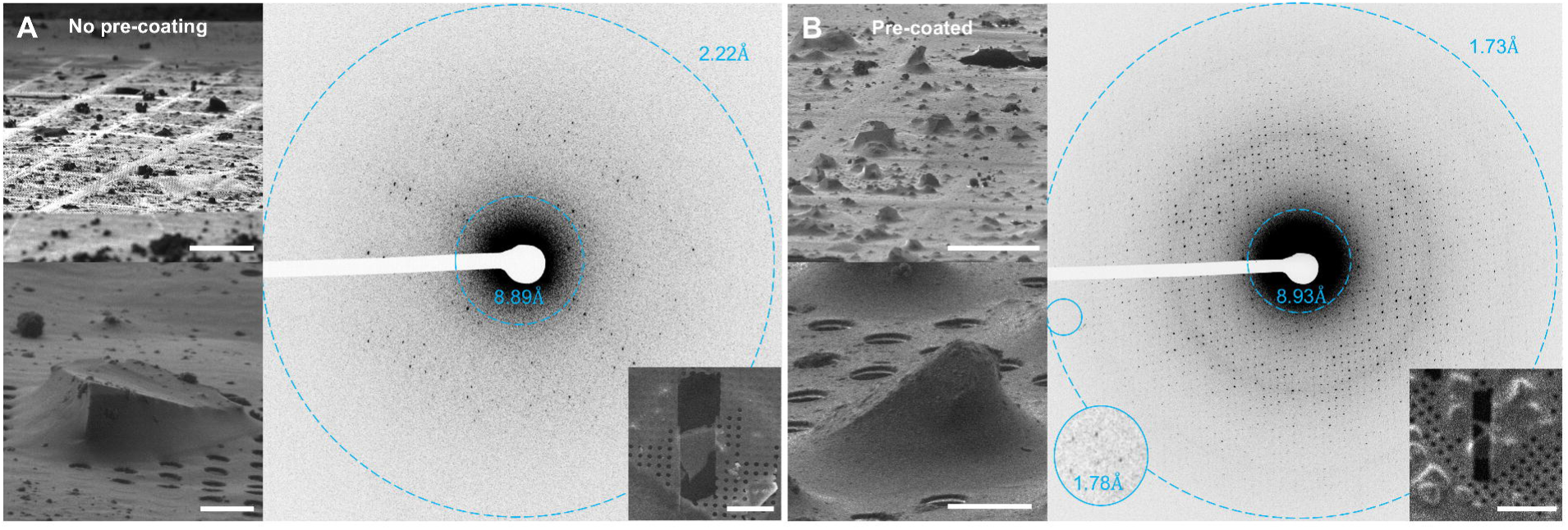
Platinum coating crystals versus no coating. FIB milling and MicroED data collection from grids without (**A**) or with (**B**) a thin layer of platinum precoating. Both were polished. Grid overviews (top left) show the difference in appearance in grids imaged by the focused ion beam. Crystals prior to milling and data collection (bottom left), and the corresponding diffraction (right) from these milled lamellae (right, inset). Scale bars 50μm top left, 5μm bottom left, 20μm right inset.

### Dose fractionation structure of Proteinase K

To compare the overall structures from both preparations with minimal TEM exposure, we merged the first half of each dataset for either the uncoated or coated lamellae that had been polished. These merged datasets were of overall high completeness, multiplicity, and were subjected to a total exposure of only 1.5e^−^Å^−2^. We refer to these as “low-dose” (LD) for the uncoated merge, and “low-dose platinum” (LDPT) (**Table 2**). The resolution for these structures was again chosen as the lowest resolution beyond which CC_1/2_ was no longer significant as indicated by student’s t-test at the p=0.1 level.

**Table 2.**
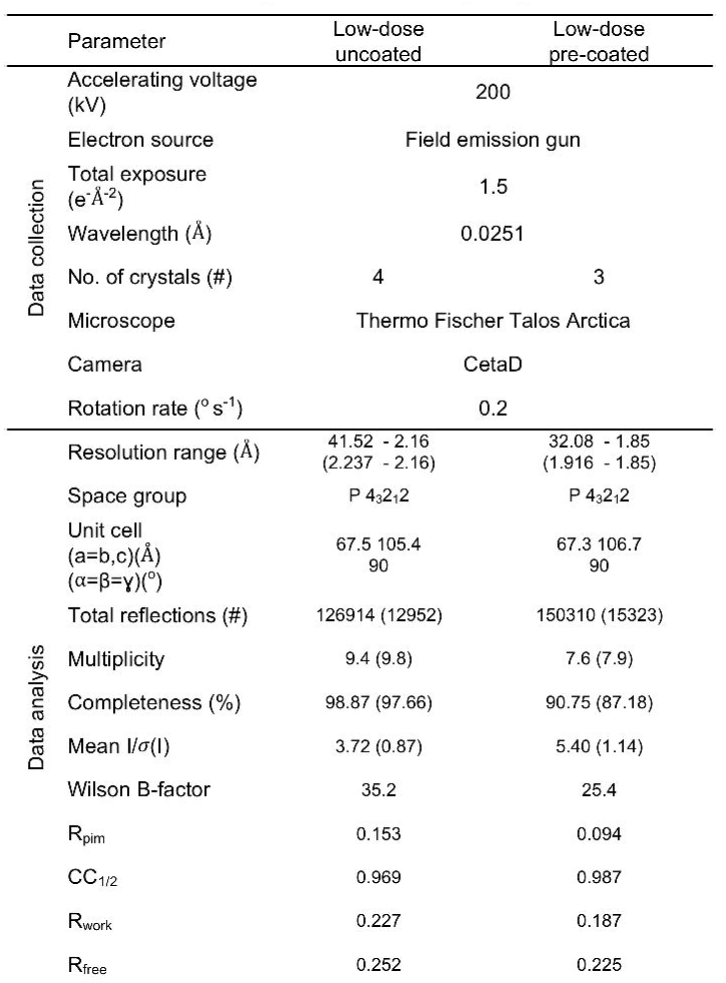
MicroED data and analysis from low-dose, merged crystals.

It appears that pre coating the samples prior to milling led to a higher resolution structure with improved refinement statistics. We found that the low-dose, uncoated dataset had a resolution of 2.16Å with an R_pim_ of 15.3%, and mean | /σ | of 3.7, whereas the pre-coated dataset extended to 1.85Å with an R_pim_ of 9.4%, and mean | / σ | of 5.4 using the same criterion. The similarities, and successful merging of these the individual crystal lamellae indicate that there were no issues with non-isomorphism between these structures. All the relevant structural and merging statistics of this structure can be found in **Table 2**.

## Discussion

The overall statistics for each set of crystals (**Figure3, Table 1**) indicate that coating with platinum may lead to higher success rate and result in higher quality data. Every lamella that produced indexable diffraction in this study resulted in a high-resolution structure solution. Only half of the uncoated crystals (4/8) could be successfully processed, whereas all (5/5) of the coated crystals yielded well-refined structures (**Figure 3**). Two crystals from each grid were milled without a final polishing step. The coated but unpolished crystal lamellae produced diffraction that ultimately led to a structure, but these lamellae had overall poorer quality than the coated, polished lamellae (**Table 1**). These results are consistent with disruption of the lattice by radiation damage, resulting in diminished resolution and thus poorer processing statistics (Hattne et al., 2018a).

**Figure 3.**
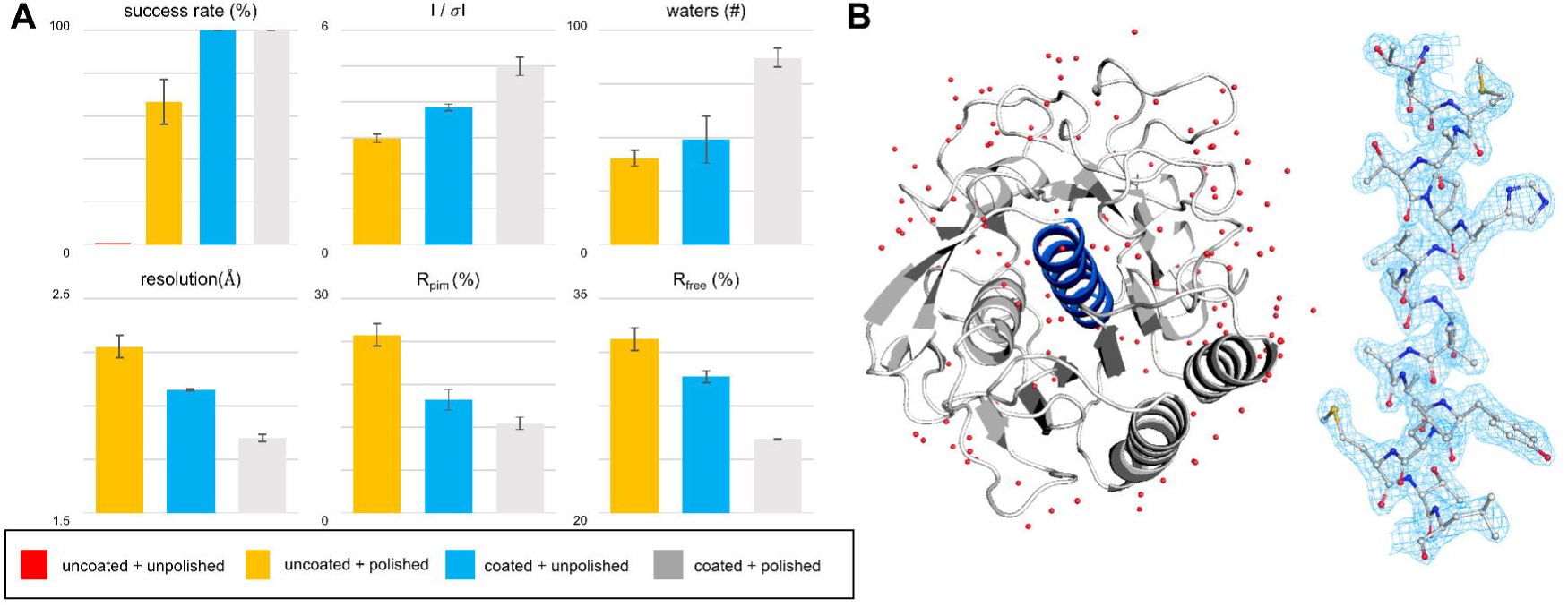
Qualitative analyses of FIB milling results and final dose fractionated structure of Proteinase K. (**A**) qualitative comparison of refined structure solutions from lamellae under different conditions. Coating the crystals resulted in 100% success rate. Crystals that were both platinum coated and polished consistently gave superior results with higher resolution, better signal-noise ratio and better refinement statistics. Error bars correspond to the standard error of the mean. (**B**). Low-dose structure of proteinase K from the platinum coated, polished datasets. The central helix corresponding to residues 328-345 is in blue with the corresponding 2mF_o_-dF_c_ density contoured at the 1.5σ level appears to the right of the structure.

The analyses indicated that pre-coating together with polishing yielded the best results. The quality metrics of success rate, maximum resolution, R_pim_, | σ | ^−1^, number of modeled waters, and R_free_ were chosen to compare between the uncoated and pre-coated lamellae (**Figure 3**). Here, differences are more pronounced between the different approaches. The uncoated and unpolished lamellae were completely unsuccessful and therefore omitted from the statistical results. The pre-coated and unpolished lamellae tended to be better than those uncoated and unpolished. The best data typically came from lamellae that were both pre-coated and polished (**Figure 3**).

The dose fractionated datasets created from both the uncoated and pre-coated, polished lamellae were of overall much higher quality than any of the individual lamellae (**Table 2**). Comparing individual statistics between the two merges clearly demonstrates that pre-coating the grids led to better quality data. The final structure and a representative portion of the 2mF_o_-dF_c_ map for the pre-coated, dose fractionated proteinase K is shown in **Figure 3**. Continuous, unbroken density was observed for disulfide bonds indicating that dose fractionation resulted in structures with minimal damage from exposure to the electron beam.

We chose the resolution limit of each dataset based on the significance of the CC_1/2_ value (p=0.1, Student’s t-test) as previously described (Evans and Murshudov, 2013; Karplus and Diederichs, 2012). Lamellae coated with platinum consistently resulted in higher resolution. Though we believe that primary gallium damage is not the culprit for the reduction of resolution, we cannot discount secondary effects that may constitute the majority of damaging events in traditional macromolecular crystallography (Garman and Weik, 2015; Hattne et al., 2018b). The coating comes with the added benefits of increased contrast during the FIB/SEM stages and prevents exposure to electrons and ions during the FIB/SEM experiments. The total exposures to both sets of crystals during MicroED experiments in the TEM were identical. The stability added by the platinum layer may serve to increase the strength of the base both while it experiences a shear force from the ion beam and during the mechanical stresses of loading and unloading the sample from the FIB/SEM to the TEM. Our initial observations of lamellae breaking during the transfer process of uncoated grids would seem to support this hypothesis.

Our platinum layer (10-50nm) is significantly thinner than the carbon-rich platinum layers used in Duyvesteyn *et al*., Li *et al*., and, later, Zhou *et al*., where a gas injection system (GIS) was used for platinum coating (Duyvesteyn et al., 2018; Li et al., 2018; Zhou et al., 2019). For example, the platinum thickness in Duyvesteyn *et al*. was estimated at 2μm thick. A thicker platinum layer may further reduce radiation damage to the specimens from the gallium beam during the milling procedure. However, thicker layers make locating smaller crystals more difficult by washing out the fine features seen with either a thin or no platinum layer, and may even not cover the areas occluded by sharp edges as pointed out in Zhou et al.(Zhou et al., 2019). Indeed, we found that even our thin layer of platinum coating made locating crystals using FIB imaging more difficult than in the uncoated grid.

All the data collected in this study was of higher resolution than our previous FIB milling investigation (Martynowycz et al., 2019). In our original report, the resolution was limited to ∼2.7Å using a total exposure of ∼4e^−^Å^−2^. Here, we used a much lower total exposure and the best resolution was 1.79Å, or about an Ångström better than our previous investigation of the same sample. Lowering the exposure by ∼1e^−^A^−2^ in total could explain some of the differences, particularly at high resolution.

We made no attempt at collecting MicroED data from crystals coated with other materials, *e.g.* silver, chromium, or carbon. Differences between coating material as well as material thickness may be important parameters in optimizing MicroED data quality. A thorough investigation of the effects of coating thickness and coating material are avenues for future investigations.

## Conclusions

We investigated the effects of pre-coating grids with platinum and polishing milled lamellae on MicroED data quality. Our results suggest that grids pre-coated with a thin layer of platinum consistently improved the quality of the MicroED data. Adding a polishing step at very low ion-beam current was necessary to observe diffraction for crystals on the uncoated grid, but did not prevent successful integration of diffraction data from pre-coated grids. We merged the first half of each dataset from either the coated or uncoated, polished grids to create complete, dose fractionated datasets with minimal exposure and better statistics for comparison. Even in this case, the very best uncoated crystals were of poorer overall quality than the pre-coated crystals. We suggest that pre-coating limits radiation damage to the crystals prior to data collection, increases the physical robustness of the preparations and leads to improved crystallographic statistics and structure. Our results set the standard for FIB milling crystals for MicroED data collection, and should ultimately result in higher resolution structures of better quality in future experiments utilizing FIB milling for sample preparation.

## Acknowledgements

The Gonen lab and the Jensenlab are supported by funds from the Howard Hughes Medical Institute. Part of the work was done in the Beckman Institute Resource Center for Transmission Electron Microscopy at Caltech. The structures presented in this investigation were deposited in the protein data bank (PDB) under the accession codes: XXXX, and in the electron microscopy data bank (EMDB) under accession codes EMD-XXXX.

## Materials and Methods

### Crystallization of Proteinase K

Proteinase K (*E. Album*) was purchased from Sigma and used without further purification. Crystals of proteinase K were grown via the sitting drop method. Proteinase K powder was dissolved in Tris-HCl 50mM pH 8.0 to a final protein concentration of 25mg mL^−1^. Sitting drops were set in microbridges in a 24-well plate by mixing 2μL of protein solution with 2μL of 1.25M Ammonium sulfate. Crystals appeared over night with high density and an average size of about 50μm.

### Sample preparation

Protein crystals were placed on glow-discharged Quantifoil R2/2 Cu200 mesh grids by pipetting 2μL of sample directly on to the carbon side. Grids were gently blotted from the back manually for 5 seconds at 100% humidity and 4°C before plunged into liquid ethane. Vitrified grids were transferred into liquid nitrogen, and clipped prior to milling and MicroED measurements.

### Ion beam milling of crystalline lamellae

All milling and coating experiments were performed using an FEI VERSA FIB-SEM instrument at liquid nitrogen temperatures. One of the grids was coated with a thin layer of platinum prior to milling and imaging using a sputter coating system built into the Quorum transfer station as previously described (Schertel et al., 2013). Crystals were identified in low-magnification SEM and FIB imaging using currents of under 100pA with 100ps dwell times. Crystals were milled by rastering over the area on interest, slowly removing portions of the embedded crystals. The current was stepped down slowly from >300pA to 30pA as the thickness of the crystals decreased to ∼300nm. For unpolished lamellae, the last 50nm on either side was removed at a beam current of 30pA to the final thickness of ∼200nm. Alternatively, to polish the lamellae, the FIB current was lowered to 10pA that was used to polish away to final 50nm above and below the initial ∼300nm lamellae to a final thickness of ∼200nm.

### Collection of MicroED data from milled crystals

Grids containing milled, crystalline lamella were loaded into the FEI Talos Arctica at liquid nitrogen temperatures. MicroED data were collected as movies. The typical MicroED workflow and setup are described in detail elsewhere (Jones et al., 2018; Martynowycz and Gonen, 2018; Martynowycz et al., 2019; Shi et al., 2016). Each crystalline lamella was continuously rotated in the electron beam at a rate of 0.2 ° sec^−1^ for a total of 60° at an exposure of 0.01e^−^ Å^−2^s^−1^ for a total exposure per crystal of 3e^−^Å^−2^. Frames were read out every 3 seconds using an FEI CetaD 4kx4k CMOS detector binned by 2. A select area aperture was used to reduce the contributions from the non-crystalline surroundings of the lamella.

### Data reduction and processing

Individual diffraction frames in MRC format from the detector were converted to SMV format for processing by in-house developed software as previously described (Hattne et al., 2015). This software is freely available on our website (https://cryoem.ucla.edu/). Diffraction images were reduced, indexed, and integrated in XDS (Kabsch, 2010a), and datasets were merged and scaled in XSCALE (Kabsch, 2010b, 2010a). Structures were determined by molecular replacement in Phaser (McCoy et al., 2007) using the search model 6CL7. The R_free_ flags were copied from this model to reduce bias (Afonine et al., 2010; Hattne et al., 2018a; McCoy et al., 2007). Models from molecular replacement were refined in Phenix.refine (Afonine et al., 2012) using electron scattering factors using automatic picking of solvent molecules (Afonine et al., 2012; Peng, 1998). No manual curation of the models was conducted in order to remove bias from the results.

